# Alpha/beta power compresses time in sub-second temporal judgments

**DOI:** 10.1101/224386

**Authors:** Tara van Viegen, Ian Charest, Ole Jensen, Ali Mazaheri

**Affiliations:** School of Phychology, University of Birmingham Birmingham B15 2TT, UK

**Keywords:** Alpha, beta, time estimation, temporal judgement, EEG

## Abstract

While the perception of time plays a crucial role in our day-to-day functioning, the underlying neural mechanism of time processing on short time scales (~1s) remains to be elucidated. Recently, the power of beta oscillations (~20 Hz) has been suggested to play an important role in temporal processing. However, the paradigms supporting this view have often had confounds of working memory, as well as motor preparation. In the current EEG study, we set out to investigate if power of oscillatory activity would be involved in time perception without an explicit working memory component or confound of a motor response. Participants indicated through a button press whether the time between a tone and a visual stimulus was 1 or 1.5s.Critically, we focused on the differences in oscillatory activity in the alpha (~10 Hz) and beta (~20 Hz) ranges preceding correct versus incorrect temporal judgments. Behaviourally, we found participants made more errors on the long (1.5s) than on the short (1s) interval. In addition, we found that participants were fastest to correctly respond to a long interval. The onset of the tone induced a suppression of alpha and beta activity over occipital and parietal electrodes. In the long estimation intervals, this suppression was greater for correct than incorrect estimations. Interestingly, alpha and beta suppression allowed us to predict whether participants would judge the long interval correctly. For the short interval trials we did not find a significant difference in alpha or beta band activity for the correct and incorrect judgments. Taken together, our behavioural and EEG results suggest a multifaceted role of alpha and beta activity in the temporal estimation of sub- and supra-second intervals, where power increases seem to lead to temporal compression. Higher alpha and beta power resulted in shorter temporal judgments for sub-second intervals.

**Highlights:**

- Temporal judgments without motor confounds were studied with EEG.
- Alpha/beta activity differences for correct and incorrect temporal judgments.
- Sub-second intervals were judged as short when alpha/beta power was higher.

## 1. Introduction

Time perception and human experience are tightly bound and play an important role in our everyday life (e.g. when playing whack-a-mole at a fun fair). Understanding how we perceive the passage of time has been an endeavour in psychology and neuroscience for over half a century (Matthews & Meck, 2016; James, 1890). And although it is well known that the human brain has a dedicated brain region for circadian rhythms (Turek, 1985), the neural underpinnings of time perception on shorter time scales (~1s) remain to be elucidated (Muller & Nobre, 2014).

Recent studies (Bartolo & Merchant, 2014; Kulashekhar et al., 2016; Kononowicz & van Rijn, 2015) suggest that beta oscillations (~15-25 Hz) play an important role in temporal processing. Kononowicz & van Rijn (2015) found that in a self-paced key-press task, trial-to-trial beta power over a central-motor electrode site positively correlated with the length of produced durations. Crucially, this correlation was present for the pre-interval period (before the first button press) and during the interval period (between the first and second button press). However, given changes of the beta rhythm locked to the onset of motor responses (Pfurtscheller & Lopes da Silva, 1999), it remains to be elucidated if beta oscillations, outside the scope of motor production, are involved in temporal judgments.

Kulashekhar et al. (2016) found that beta band power increased while storing and retrieving temporal but not colour information. This important finding suggests that beta oscillations are indeed involved in temporal judgments, specifically in the memory encoding and retrieval of temporal information. However, it remains unclear if beta oscillations were directly involved in the perception of time, because temporal information was contrasted with colour information.

In the current study, we set out to investigate how the modulations of oscillatory activity in the alpha (8-12 Hz) and beta range would be involved in a temporal perception task without an explicit working memory component or confound of a motor response. In addition to activity in the beta range, we focused on alpha activity (~10 Hz) given its previously established role in working memory (Lakatos et al., 2008; Haegens et al., 2010; Wilsch et al., 2014), cross-modal attention (Van Diepen and Mazaheri, 2017, Foxe et al., 1998), and perception (Van Dijk et al., 2008). We utilized a simple forced-choice temporal estimation task where participants indicated by button press whether they judged the time between a tone and a visual stimulus as 1 or 1.5s. We focused on the differences in EEG activity between correct and incorrect temporal judgments in the interval between the tone and visual stimulus. In addition to looking at the condition differences in alpha and beta power between correct and incorrect judgments, we also examined if on a trial-by-trial basis power modulations in those frequencies predicted accuracy of temporal judgments.

## 2 Materials And Methods

### 2.1 Participants

Twenty-three healthy, young adults (right and left handed) participated in this study. Data from three participants was excluded (one participant presented severe muscle artefacts (>25% of trials), one participant did not make any errors in the short interval, one participant made only 5 errors in the long interval). Twenty datasets were used for further analyses (13 females, mean age: 27 years, range: 19-41 years, 3 left handed). Participants had normal or corrected-to-normal vision and hearing and they had no known history of neurological disorders. In concordance with university guidelines, participants were paid £6 or 1 participation credit per hour. Participants gave written informed consent before data collection. Ethical approval was provided by the Ethics Committee of the University of Birmingham.

### 2.2 Paradigm

We utilized a novel two-forced-choice temporal judgement paradigm where the participant’s task was to estimate whether the time between an auditory stimulus and a visual stimulus was short (1s) or long (1.5s), via a button press (Fig. 1). The paradigm was programmed in Matlab (Natick, MA) using Psychophysics Toolbox extension (Brainard, 1997; Pelli, 1997; Kleiner et al, 2007; psychtoolbox.org). The auditory stimulus was a pure tone of 1000 Hz, which was administered through headphones (Sennheiser, HD 280 PRO) and lasted for 50ms (including a 5ms rise and fall shaped by a Blackman window). The tone signified the start of the interval, which was followed by a visual stimulus that indicated the end of the interval. The visual stimulus was a Gabor patch (angle 5° clockwise, contrast 80%, spatial frequency 10 Hz, phase 0°), which lasted for 50ms. Short and long intervals were randomly interspersed.

**Figure 1.**
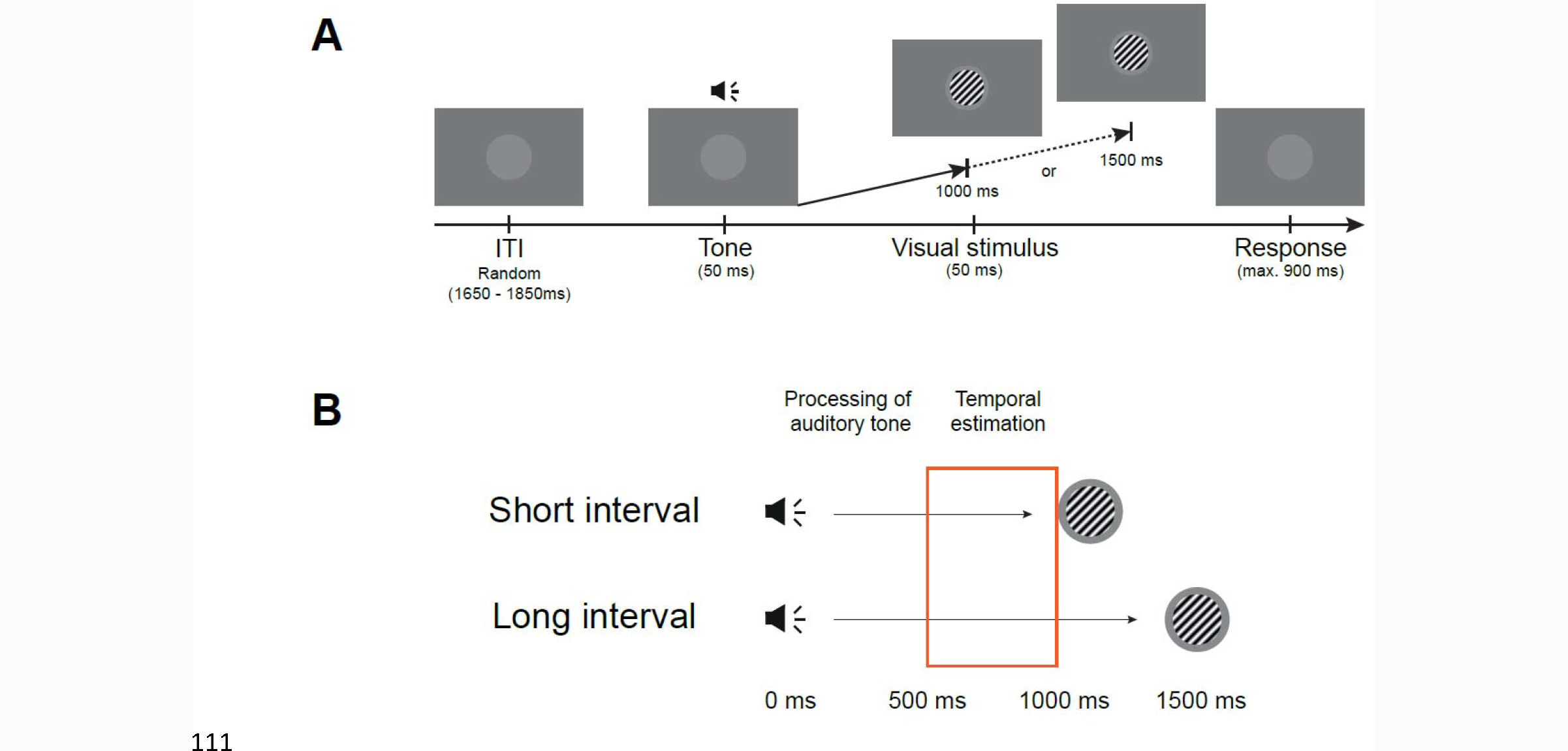
Experimental paradigm and predictions. Participants performed a two-forced-choice task estimating the time interval between the tone and the visual stimulus (A). Trials were initiated with the presentation of a 1000 Hz pure tone for 50 ms. The tone was followed by a Gabor patch, which lasted for 50ms, 1 or 1.5s after the tone was presented. Participants judged whether the interval between the tone and the visual stimulus was short (1s) or long (1.5s) by pressing a button. Participants had 900ms to respond. After the response (or 900ms) a new trial was initiated. A light grey placeholder was always on the screen to minimize eye movements. ITI = inter-trial interval. Our time window of interest was before the short interval was over (1000ms) and after processing of the auditory tone (~500ms; B). We hypothesized that differences between correct and incorrect trials would be caused by differences in time estimation if the difference lay in the time window indicated by the orange box.

After the visual stimulus participants responded with a button press. If participants did not respond within 900ms, a screen was displayed that read “Too slow! Please respond faster.” Participants were pushed to respond fast to enhance the likelihood that they would make errors. Participants used the index (left button) and middle (right button) finger of their right hand to respond. Response buttons were randomly counterbalanced across participants. The dataset presented here contains data of 11 participants that pressed the left mouse button for the long interval and 9 participants that pressed the right mouse button for the long interval.

Participants were seated at a comfortable distance from the computer screen. The distance was then measured and used to ensure a visual angle ~4° across all participants. To assure participants kept their eyes on the screen a light grey placeholder was presented centrally on the grey background (Fig. 1). The placeholder was 5% larger than the Gabor patch, so the Gabor patch fell inside the placeholder.

Participants practiced for 3 blocks that consisted of 12 trials each. Participants received feedback after each block. The EEG recording started by collecting resting state data with eyes open and eyes closed for 30s each. This procedure was performed twice, randomly starting with either eyes closed or eyes open data collection. The resting state EEG data was not used for the purpose of this study. Following this, participants performed 16 blocks of 60 trials, yielding a total of 960 trials per participant. Each block lasted ~3min and participants were instructed to rest their eyes in between blocks. The refresh rate of the screen was 60Hz and all timings of the experiment were set as multiples of this refresh rate.

### 2.3 Behaviour

Behavioural measures consisted of response times and error rates. Response times were calculated from visual stimulus onset until response. Errors were calculated for each interval condition separately by calculating the percentage of errors (i.e.judgment accuracy) relative to the total number of trials in that condition. Statistical analyses were performed on response times and errors, where the subject-specific averages were subjected to repeated-measures ANOVA. For response time the factors were interval (short or long) and response (correct or incorrect). For judgement accuracy (i.e. correct and incorrect judgments) the factor was interval (short or long).

### 2.4 EEG Data Acquisition

EEG was acquired using the EEGO Sports system (ANT Neuro, Enschede, Netherlands) and Waveguard caps housing 64 Ag/AgCl electrodes arranged in a 10/10 system layout (including left and right mastoids, CPz as reference and AFz as ground). Impedances were kept below 20kΩ, and the data was acquired using a sampling rate of 500Hz. EOG was collected for horizontal eye movements, by placing bipolar electrodes on the outer canthi of the left and right eye. ECG was collected for heart rate data, by placing one bipolar electrode on the right chest, one bipolar electrode on the left abdomen and one bipolar electrode on the left collar bone. This latter electrode acted as a ground electrode for the ECG signal. Offline data analysis took place in Matlab with eeglab functions (version 13.1.1b; Delorme & Makeig, 2004) and the FieldTrip software package (Oostenveld, Fries, Maris & Schoffelen, 2011).

### 2.5 Preprocessing

#### 2.5.1 TFR analyses

Using EEGLAB, the data was epoched around −1000 to 3500 ms from tone onset. Subsequently, a baseline from −100 to 0 ms was applied to the data to reduce DC offset. Missed trials were discarded, as well as trials where the timing of the event triggers deviated more than 2 ms of the intended timing. This last step was performed because timing was crucial in this experiment and we wanted to exclude trials that deviated more due to unforeseen slowness of the operating system of the stimulus computer. Based on visual inspection trials were removed for the following reasons: muscle artefacts, noise (i.e. electrode jumps or other electrode related noise), horizontal eye movements and blinks at visual stimulus presentation. An average reference (excluding bipolar electrodes) was applied to the cleaned data, and then CPz was reconstructed. Ocular artefacts were removed in FieldTrip using independent component analysis (ICA, infomax algorithm) incorporated as the default “runica” function. Prior to the ICA, a PCA (15 components) was performed on the data, to speed up the ICA procedure.

#### 2.5.2 Time Frequency Representation

Using the FieldTrip function ‘ft_freqanalysis_mtmconv’ time frequency representation (TFR) of power was obtained for each trial by performing a fast fourier transform using a Hanning taper in combination with a sliding time window. The time window was adapted to the frequency of interest (Δ *T* = 3/*f*) The frequency range of interest was from 2 to 40 Hz in steps of 1 Hz. TFRs were calculated for the long and short interval and for correct and incorrect responses separately, leading to 4 different subsets of trials. After we assessed that there were no differences in baseline (i.e. pre-cue) oscillatory power for our frequency bands of interest between correct and incorrect responses, data in each condition was normalized to be the relative change in power according to the following formula,

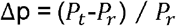

where *P_r_* was the mean power during the pre-cue period (700 – 200ms before tone onset) and *P_t_* was the power at each specific time point.

### 2.6 2.6 Statistics

#### 2.6.1 TFR analyses

Our selection of frequency bands were loosely based on previous literature (Palva & Palva, 2007; Weisz et al., 2011; Zumer et al., 2014). Our statistical comparisons were made for the alpha (8-14 Hz) and beta (15-25 Hz) band, between correct and incorrect responses for the short and long interval separately. We did not directly compare the short and long intervals, due to the fact that the amount of spectral leakage from the response evoked by the auditory tones was different over short and long intervals.

The differences in oscillatory power between conditions were statistically assessed by means of the cluster level (channels and time-points) randomization approach (Maris and Oostenveld, 2007). Here, the power of the frequencies of interest in each channel and time point within the time intervals of interest, were clustered according to exceeding a threshold of *p* < .05 obtained from a two tailed dependent samples t-test. The time interval of interest was 500-1000ms after tone onset for both frequency bands (Fig. 1B). Note that this was well after processing of the auditory cue or start point of the interval and when the short time interval ended. Next, the Monte Carlo p-values of each cluster were obtained by randomly swapping the condition labels within participants 2500 times. A difference between conditions was deemed significant if the cluster *p*-value was smaller than .025 (two-sided test).

#### 2.6.2 Power differences predictive of temporal judgments accuracy

Finally, we asked whether power differences in the clusters we identified were predictive of time-estimation accuracy. To assess this, at the first level we fitted logistic regression models (with the Matlab function ‘glmfit’) of alpha or beta power, averaged over the cluster that showed the maximal difference between conditions, as a predictor of response accuracy (short or long). To apply this model to ordinal data we used the logit link function. This model yields a Beta weight for each participant. We then tested the significance of these Beta weights across-participants with a permutation test. In this permutation test a random sign was assigned to the observed Beta values, which allowed us to build a distribution for the null hypothesis. We then checked whether our mean observed Beta value was as extreme or more extreme than the outer 2.5% of the null distribution (two-sided test).

To visualize our effect, we approached our data in two ways. First, we plotted accuracy as a function of binned z-scored alpha and beta power in 5 equal-sized bins within a participant. We used z-scored power because this reduced the variability between participants. The z-scored power was calculated according to the following function:

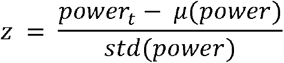

where z is the z-scored power, power_t_ is the power for the trial on which the z-scored power is calculated, μ(power) is the average power of all trials and std(power) is the standard deviation of all trials. Second, we plotted the logistic regression lines for normalized power within participants. Normalized power was obtained as following:

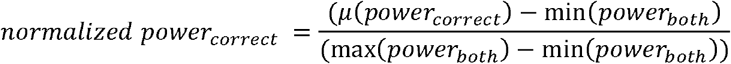

where normalized power_correct_ represents the normalized power for each trial for correct trials. Power_both_ refers to power of both correct and incorrect responses. This approach yielded normalized power values between 0 and 1, which ensured an equal length for each of the regression lines. Please note that the regression line is the same for normalized or unnormalized power. Moreover, the statistics explained above were only applied to the Beta values that were obtained from logistic regression with baseline corrected power data.

## 3. Results

### 3.1 Behavioural

#### 3.1.1 Participants respond fastest on long correct trials

We first investigated if there were response time differences between correct and incorrect responses on the short and long interval. In terms of main effects, we found a significant effect of response (*F*(1,19) = 49.882, *p* < .0001, η_p_^2^= .724), with responses being fastest for correct responses (384.2 vs. 456.9ms). In addition we found a significant effect of interval and interval length (*F*(1,19) = 138.477, *p* < .0001, η_p_^2^ = .879), with response time being fastest for longer (i.e. 1.5s) versus shorter (i.e. 1.0s) temporal judgments (395.9 vs. 447.2ms).

We found that participants were fastest to respond in correctly perceived long intervals, with a significant interaction effect of response × interval length (Fig. 2A; *F*(1,19) = 33.394, *p* < .0001, η_p_^2^= .637). Post-hoc analyses revealed for the longer temporal judgments individuals were significantly faster on correct responses (329.0 ± 47.7ms, mean ± standard deviation) than incorrect responses (458.8 ± 66.9ms). However, on the short interval we observed no significant difference in response time between correct (439.3 ± 61.3ms) and incorrect (454.9 ± 81.2ms) responses.

#### 3.1.2 Participants made more errors during the long interval

We found that individuals made significantly more errors during the long interval trials than during the short interval trials (long interval errors: 25.91 ± 13.23%; short interval errors: 9.68 ± 6.54%; *F*(1,19) = 19.893, *p* ≈ .0003, η_p_^2^ = .511).

**Figure 2.**
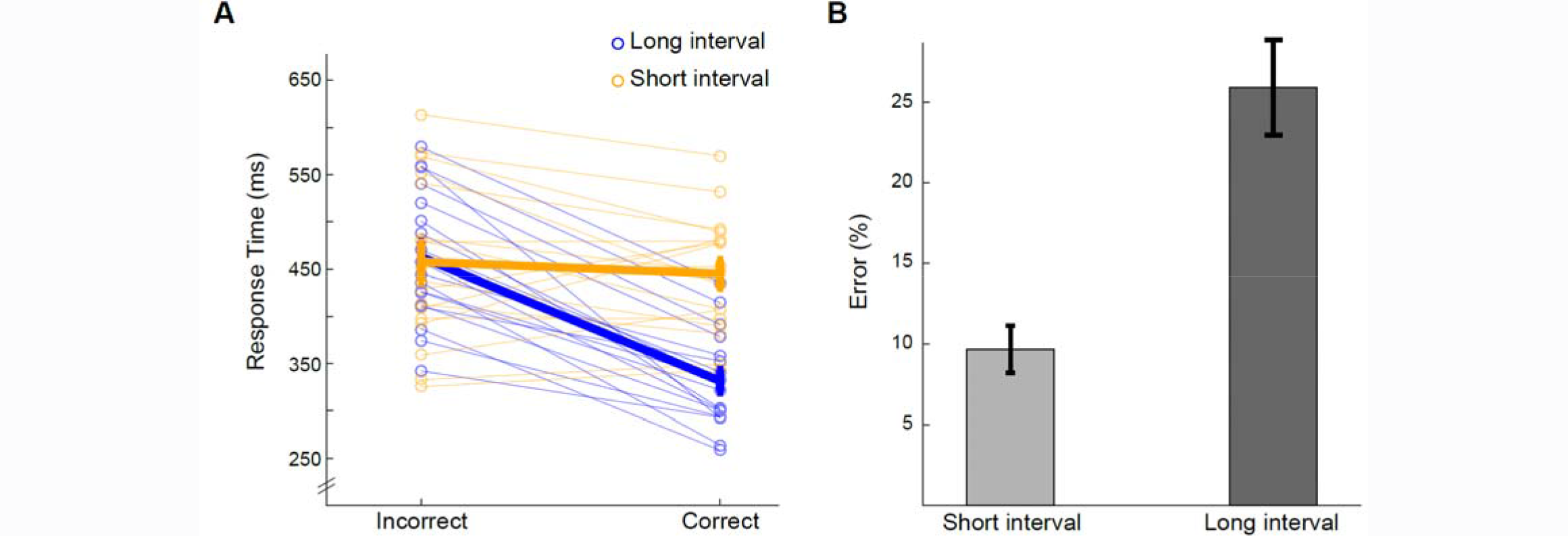
Behavioural effects. On the long interval (A; blue lines), individuals were significantly faster on correct responses (329.0 ± 47.7ms, mean ± standard deviation) than incorrect responses (458.8 ± 66.9ms). On the short interval (orange lines), no significant difference in response time between correct (439.3 ± 61.3ms) and incorrect (454.9 ± 81.2ms) responses was found. Individual participant data is represented in opaque lines with open dots, participant average data is depicted in the fat lines. Individuals made significantly more errors in the long interval (B; 25.91 ± 13.23%) compared to the short interval (9.68 ± 6.54%). Error bars represent standard errors of the mean.

### 3.2 EEG results

#### 3.2.1 No difference in alpha/beta suppression in correct versus incorrect short interval (1s) judgment trials

We set out to investigate if oscillatory activity between the tone and a visual target occurring 1s later could be predictive of states in which participants were more likely to over-estimate the durations. The time-frequency representations of power between the auditory tone and visual target interval collapsed across all electrodes can be seen in Figure 3A for correct (left) and incorrect trials (right). For both correct and incorrect short interval judgments, the onset of the cue induced a transient increase in theta activity, followed by a sustained power decrease in alpha and beta bands starting ~500ms after tone onset. Shortly after the visual cue was presented a more pronounced transient decrease in beta power can be seen ~1300ms after tone onset, which roughly coincides with the button press (average RT for correct and incorrect judgments was ~447ms).

To assess whether alpha and beta suppression were significantly different between correct and incorrect judgments of the short interval, we used cluster based statistics correcting for multiple comparisons. For the alpha band we found no significant difference in the time window of interest (500-1000ms after tone onset). Subsequent examination of the alpha activity within that time found the greatest difference in alpha modulation between correct and incorrect trials to be from 750-800ms after tone onset, where correct trials had more power than incorrect trials. However, this difference was not significant (*p* > .06). Figure 3C (top) shows the average power difference between correct and incorrect trials averaged over the entire time window of interest, irrespective of the cluster.

Similarly, for the beta band we found no significant difference in our time window of interest. The difference between correct and incorrect trials was again strongest from 750-800ms after tone onset, where correct trials had more power than incorrect trials (*p* > .045, remember that our critical p-value is .025). Figure 3C (bottom) shows the average power difference between correct and incorrect trials averaged over the entire time window of interest.

**Figure 3.**
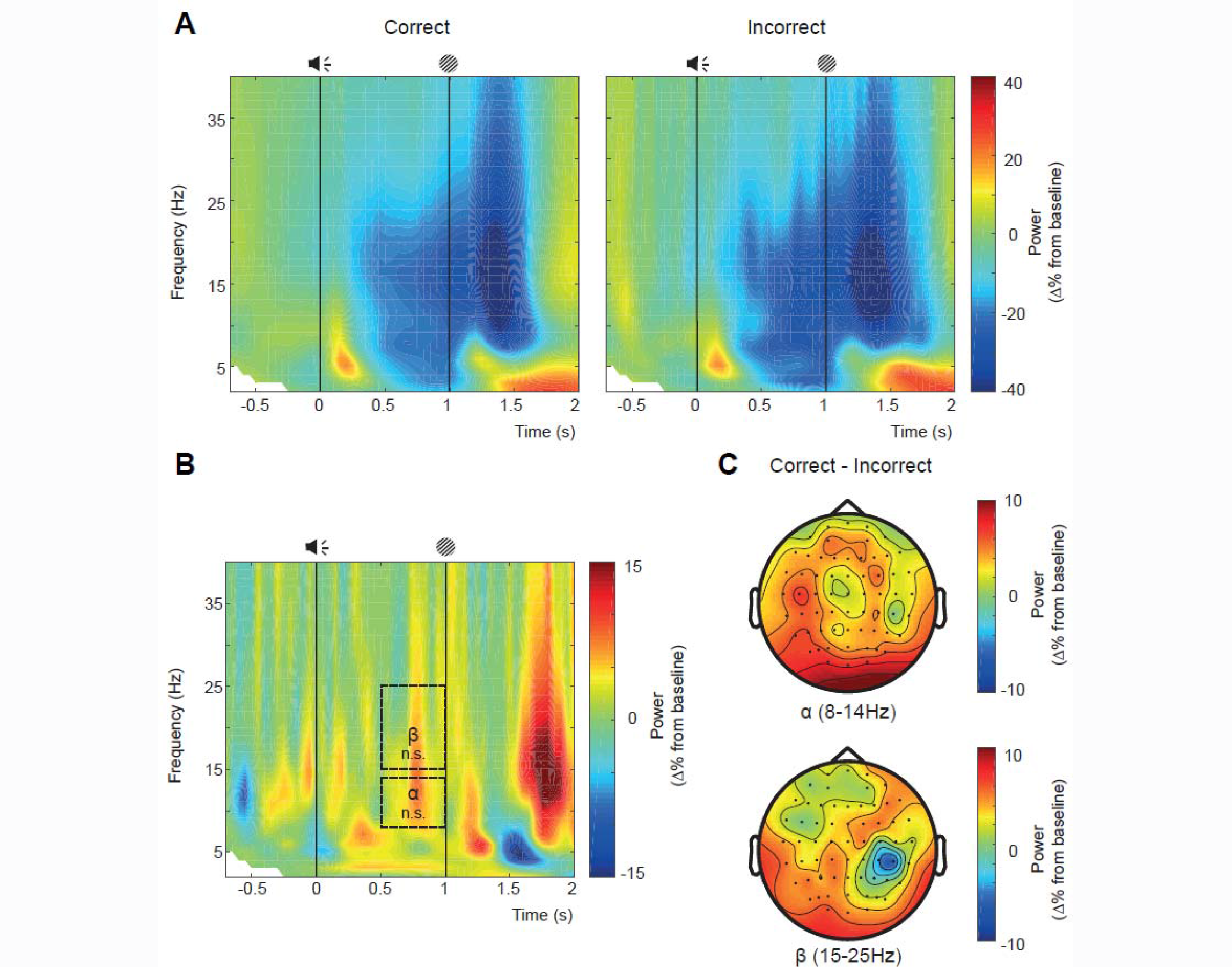
TFR for the short interval. The percentage change from baseline is plotted for the different frequency bands over time averaged across all electrodes, with a baseline interval from -700 to -200 ms, tone onset at t = 0 s followed by visual stimulus presentation at t = 1.0 s. Correctly perceived short intervals are visible on the left and trials incorrectly perceived as long are visible on the right (A). The difference plot shows more power for trials correctly perceived as short than trials incorrectly perceived as long (B). There were no significant differences observed in the alpha or beta band. The topography of the average activity over the time window of interest is plotted in C (correct – incorrect trials).

#### 3.2.2 Greater alpha/beta suppression in correct versus incorrect long interval (1.5s) judgment trials

As with the short-interval, the time-frequency representations of power between the auditory cue and visual target interval collapsed across all electrodes can be seen in Figure 4A for correct (left) and incorrect (right) trials. After the transient theta increase induced by the onset of the auditory cues, we observed a sustained power decrease in alpha and beta bands, from ~500ms to ~1700ms after tone onset. Moreover, a more pronounced transient beta decrease was observed at ~1700-1900ms after tone onset. This beta decrease roughly coincides with the button press (averaged RT for correct and incorrect trials was ~393ms).

To examine the sustained alpha and beta power decreases, we tested the interval of 500ms after tone onset until visual stimulus onset of the short interval at 1000ms. However, we should note here again that the onset of the visual stimulus was 1500ms in long interval trials.

To assess whether alpha and beta suppression were significantly different between correct and incorrect judgments of the short interval, we used cluster based statistics correcting for multiple comparisons. For the alpha band we found a significant difference in our entire time window of interest (500-1000ms after tone onset), where correct trials had less power than incorrect trials (*p* < .0024). Figure 4C (top) shows the average power difference between correct and incorrect trials averaged over the cluster where the difference was most pronounced (outlined in Fig. 4B). The black dots with white edges represent electrodes that were part of the cluster at any one time point from 500-1000ms.

Similarly, for the beta band we found a significant difference power between correct and incorrect judgments in our time window of interest. The difference between correct and incorrect trials was strongest from 600-1000ms after tone onset, where correct trials had less power than incorrect trials *(p ≈* .0004). Figure 4C (bottom) shows the average power difference between correct and incorrect trials averaged over the cluster where the difference was most pronounced (outlined in Fig. 4B). The black dots with white edges represent electrodes that were part of the cluster at any one time point from 600-1000ms.

**Figure 4.**
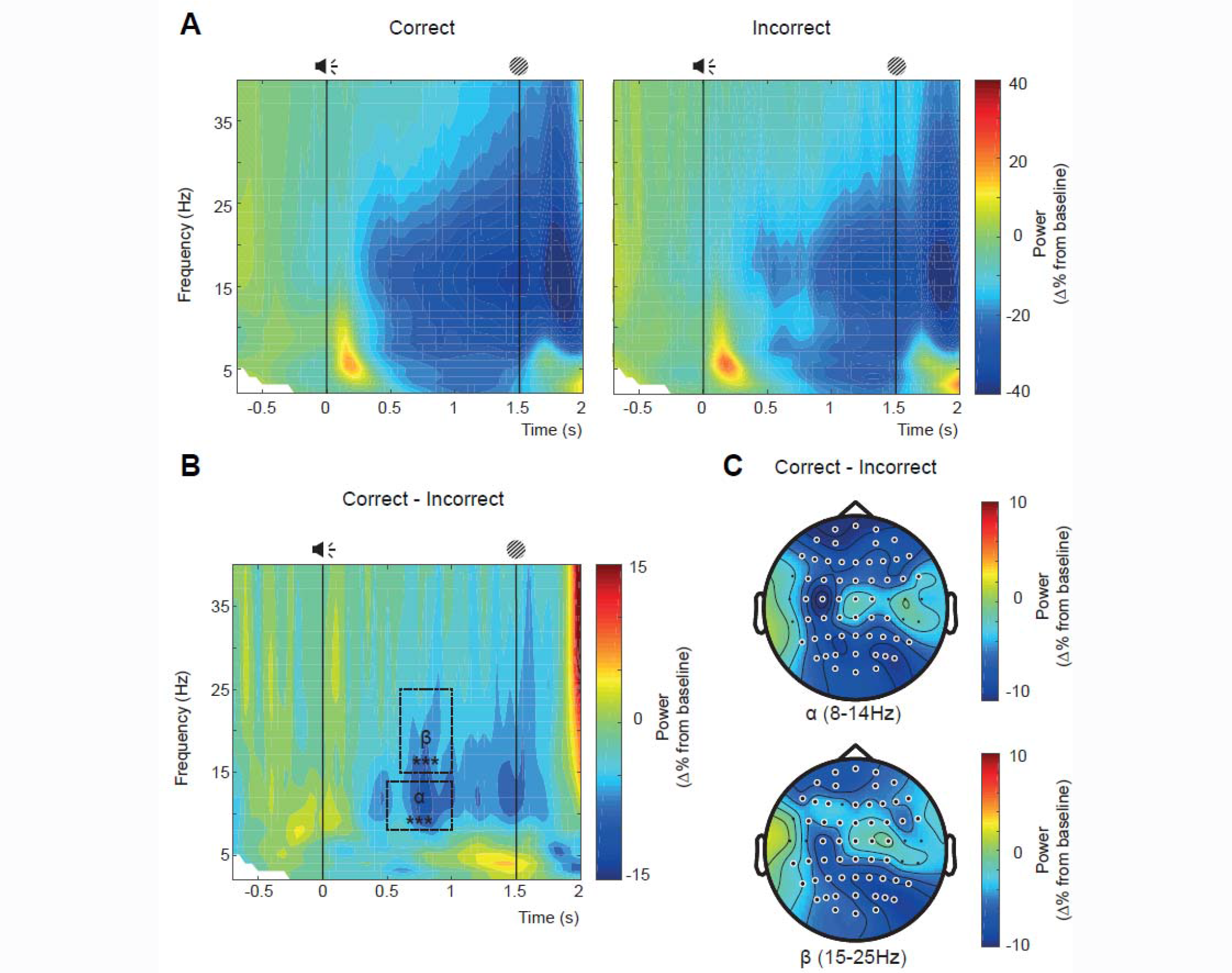
TFR and topographical distribution for the long interval. Same as in Fig. 3A but now with visual stimulus presentation at t = 1.5 s. The difference between correct (left in A) and incorrect (right in A) trial types is visualized in B, where power was lower for correct trials. Significant differences between correct and incorrect responses were observed in the alpha and beta bands. The topographical plots show correct-incorrect responses (C). Black circles with white edges represent electrodes showing significant modulation at any one time point, corrected for multiple comparisons using cluster randomization routine (Maris and Oostenveld, 2007).

### 3.3 Effect size

Finally, we calculated the effect size of our smallest TFR effect to ease future, direct replications of our reported effects. Please note that partial Eta squared was reported as effect size for our behavioural findings. To assess the effect size of our result, we used the alpha cluster depicted in Figure 4C and took the difference between the averaged power over trials for correct and incorrect responses to the long interval trials. The mean difference of these power values was -7.02% with a standard deviation of 2.18%, which leads to a Cohen’s *d ≈* 3.22. To replicate this finding a follow-up study, with 80% power and α = .05, would need 4 participants.

### 3.4 Response to auditory cue not different between correct and incorrect temporal judgments

Previous research suggested that expectation could allocate resources to a task-relevant region (e.g. visual system) at the expense of processing in the task-irrelevant regions (Cartwright-Finch and Lavie, 2007). Moreover, we wanted to make sure that any differences we observed were not caused by differences in processing of the auditory cue. As such, we investigated if the response to the auditory cue could be predictive of the accuracy of temporal judgments. However, we did not find a significant difference in power between correct and incorrect responses to the processing of the tone (50-300 ms after tone onset, 3-6 Hz). There was no cluster for the long interval. For the short interval a non-significant cluster was identified (*p* < .1), with the largest difference from 250-300ms after tone onset.

### 3.5 Alpha and beta power predict temporal judgments in the long interval

Here, we investigated whether a relationship between correctly identifying the temporal durations and the power of alpha and beta activity could be observed on a trial-by-trial basis To this end, we fitted logistic regressions to each individual participant’s data and tested the beta weights of these fits at the group-level against a null distribution obtained through a sign permutation test (see Materials & Methods for more details).

A logistic regression model was fitted with a logit link function where alpha power for each trial, averaged over the electrodes depicted in Figure 4C (top) and over time 500-1000ms after tone onset, was used as a predictor of correct or incorrect responses. This approach yielded a Beta value for each participant. To assess whether these Beta values were significantly different from 0 (no predictive value) we used a permutation test. We found that alpha power of the cluster we observed significantly predicted correct versus incorrect responses (*p* < .0001). Similarly, we found that beta power for each trial, averaged over the electrodes depicted in Figure 4C (bottom) and over time 600-1000ms after tone onset, predicted responses (p = .0001).

**Figure 5.**
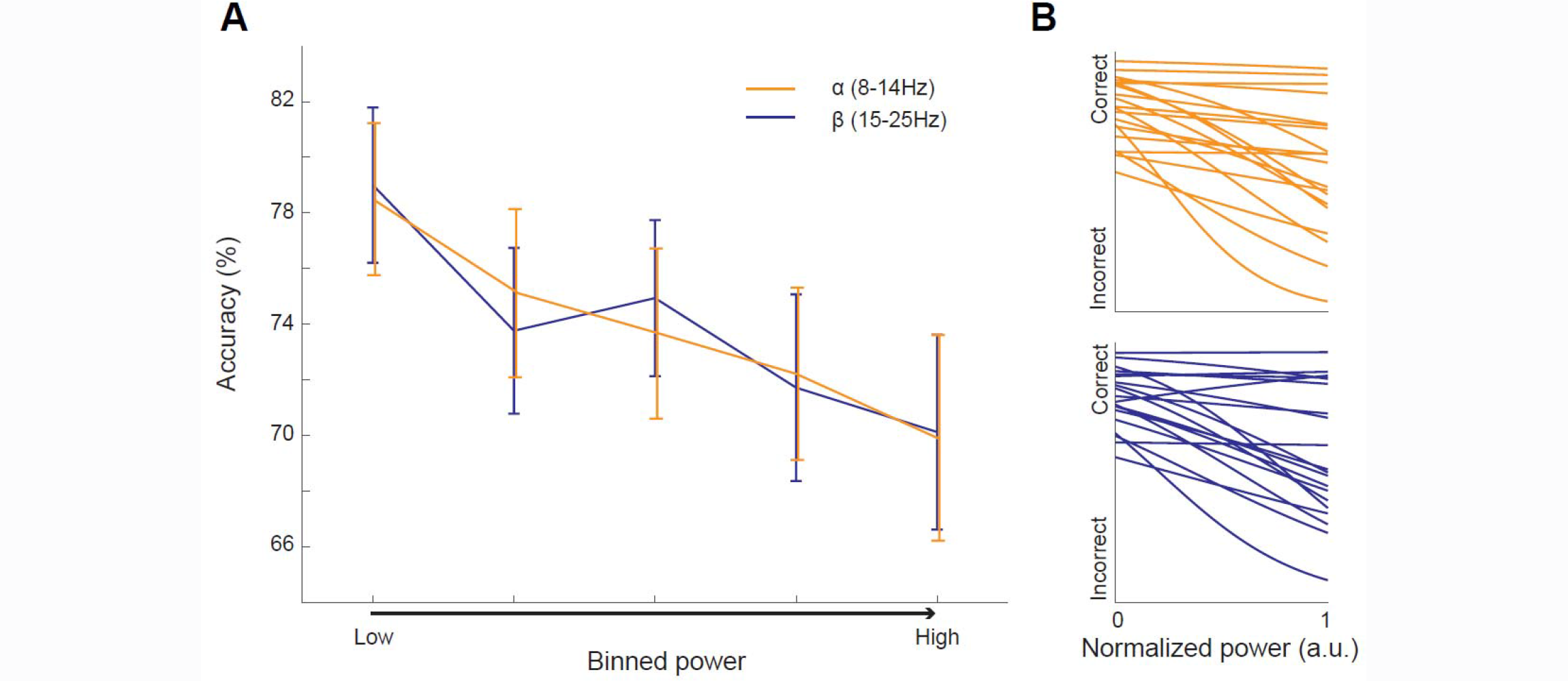
Accuracy as a function of power. Alpha and beta power from the clusters displayed in Figure 4C significantly predicted responses. This relationship is visualized here by plotting accuracy as a function of binned z-scored alpha and beta power, where each bin has the same amount of trials within a participant (A). As alpha and beta power increase, the accuracy decreases. Error bars represent standard error of the mean. The logistic regression lines for normalized power are shown in B (alpha on the top and beta on the bottom). Statistics reported in the text were performed on non-normalized power values to make sure that our normalization procedure did not drive the effect.

To further visualize this effect we binned z-scored alpha and beta power in 5 equal sized bins for each participant (same number of trials in each bin within a participant) and plotted accuracy as a function of binned power (Fig. 5A). As alpha/beta power increases participants are more likely to judge a 1.5s interval as short. We also plotted the regression lines of normalized power in Figure 5B. The power was normalized between 0 and 1 such that the regression lines were of the same length. In the supplementary materials the individual participant data can be viewed, with the logistic regression fits for each participant (Supplementary Fig. 1). The Beta values of the regression lines in the supplementary figures depicts the data on which statistics were performed.

## 4. Discussion

In the current EEG study, we set out to investigate how the power of ongoing oscillations could be involved in time perception, controlling for the presence of an explicit working memory component or confound of a motor response. Using a novel paradigm, we asked our participants to judge the temporal interval between a tone and a visual stimulus as being short (i.e 1s) or long (i.e. 1.5s). Our analyses focused on the activity preceding correct versus incorrect temporal judgments. For both correct and incorrect short interval judgments, we observed that the onset of the cue induced a transient increase in theta activity, followed by a sustained power decrease in alpha and beta bands starting ~500ms after tone onset and continuing till well after the visual stimulus. While we did not find any significant differences in alpha or beta power modulation in correct versus incorrectly judged short intervals, we did so for the long-interval judgments. Specifically, correct judgments of the long interval had significantly greater alpha and beta suppression than incorrect judgments. On a trial-by-trial basis we found that the more alpha/beta power increased in the 0.5-1s window after cue onset, the more likely the participants were to judge a 1.5s interval as short.

Behaviourally, we did not find any significant difference between correct and incorrect responses for the short interval. In contrast, for the long interval, individuals were significantly faster on correct responses than incorrect responses. Moreover, participants made more errors when estimating long duration trials (1.5s) than when estimating short duration trials (1s). This is in line with Webber’s law, which predicts that it becomes increasingly more difficult to estimate time as more time has passed by. In addition, we found that individuals responded faster on correctly observed long intervals. This may suggest that participants have a template of 1s in mind and when this time interval has passed they expect the upcoming visual stimulus at 1.5s allowing the speeded response.

Taken together these behavioural findings may indicate that two different mechanisms operate for short and long intervals. A differentiation between sub- and supra-second intervals would be in line with previous literature (Lewis & Miall, 2003; Matell & Meck, 2004), although it is disputable whether a 1s interval would be classified as a sub-second interval. Alternatively, another explanation for these behavioural differences might be that the same mechanism underlies both intervals, but that this mechanism does not get enough time to unfold in the short interval. However, this alternative explanation cannot account for our finding that individuals make more errors in the long.

Our EEG data showed no significant difference in power between correct and incorrect judgments of short interval trials.For the long interval trials, we found a significant difference between correct and incorrect responses in the alpha and beta band. Moreover, the cluster that exhibited the largest z-scored alpha difference between these trial types allowed us to predict whether participants judged a trial as short or long. Interestingly, we were able to predict the response before the short interval time window closed (<1000ms).

It remains to be elucidated why we did not find an effect of alpha/beta power for correct and incorrect judgments of the short interval. One explanation might be that we simply did not have enough incorrect judgments for these trial types, hampering our power to detect a small effect. This explanation is in line with our insignificant results, where the clusters showed more power for correct trials than incorrect trials, suggesting that power increases condense temporal judgments and bias participants to judge an interval as short. Alternatively, a different mechanism might operate for sub- and supra-second time interval judgments. Future research should investigate this.

In accordance with Kononowicz & van Rijn (2015), we found that alpha/beta power fluctuations might represent a neural signature that is used to track time. Kononowicz & van Rijn (2015) showed that self-paced intervals that were longer than the target interval had more beta power. In the current study, we showed that increased alpha/beta power leads to underestimation of the time that has passed by. Taken together, this suggests that high alpha/beta power leads to compression of the time that has passed by. From this framework it would follow that crossing a certain threshold leads to experiencing a temporal window as shorter. It will be exciting to see if these findings replicate and how the difference between sub- and supra-second time intervals can be explained, but our findings capitalize on the importance of oscillatory activity in time processing. In summary, in the current study we found that modulations in alpha and beta power were predictive of temporal judgments. Specifically, increased alpha/beta power between the cue-target interval biased participants to report the long intervals as being short. We hypothesize that fluctuations in alpha/beta power condense the subjective experience of time passing by.

## Acknowledgements

We would like to thank André Cravo for his valuable suggestion to include the analysis where we predicted responses based on power.

## Funding

This research did not receive any specific grant from funding agencies in the public, commercial, or not-for-profit sectors.

## References

Bartolo, R., Prado, L. and Merchant, H. 2014. “Information Processing in the Primate Basal Ganglia during Sensory-Guided and Internally Driven Rhythmic Tapping.” The Journal of Neuroscience: The Official Journal of the Society for Neuroscience 34 (11): 3910–23.

Brainard, D. H. 1997. “The psychophysics toolbox.” Spatial Vision, 10, 433–436.

Cartwright-Finch, U., and Lavie, N. 2007. “The Role of Perceptual Load in Inattentional Blindness.” Cognition 102 (3): 321–40.

Delorme, A., and Makeig, S. 2004. “EEGLAB: An Open Source Toolbox for Analysis of Single-Trial EEG Dynamics Including Independent Component Analysis.” Journal of Neuroscience Methods 134 (1): 9–21.

van Diepen, R.M., and Mazaheri, A. 2017. “Cross-Sensory Modulation of Alpha Oscillatory Activity: Suppression, Idling, and Default Resource Allocation.” The European Journal of Neuroscience 45 (11): 1431–38.

van Dijk, H., Schoffelen, J.-M., Oostenveld, R., and Jensen, O. 2008. “Prestimulus Oscillatory Activity in the Alpha Band Predicts Visual Discrimination Ability.” The Journal of Neuroscience: The Official Journal of the Society for Neuroscience 28 (8): 1816–23.

Donner, T.H., Siegel, M., Fries, P., and Engel, A.K. 2009. “Buildup of Choice-Predictive Activity in Human Motor Cortex during Perceptual Decision Making.” Current Biology: CB 19 (18): 1581–85.

Foxe, J.J., Simpson, G.V., and Ahlfors, S.P. 1998. “Parieto-occipital ~10 Hz activity reflects anticipatory state of visual attention mechanisms.” Neuroreport 9, 3929–3933.

Haegens, S., Osipova, D., Oostenveld, R., and Jensen, O. 2010. “Somatosensory Working Memory Performance in Humans Depends on Both Engagement and Disengagement of Regions in a Distributed Network.” Human Brain Mapping 31 (1): 26–35.

James, W. 1890. “The Principles of.” Psychology 2. http://coursecontent.learn21.org/ENG3x-HS-U10/a/unit04/resources/docs/E3HS4.COrgTablePsychology.pdf.

Kelly, S. P., and O’Connell, R.G. 2013. “Internal and External Influences on the Rate of Sensory Evidence Accumulation in the Human Brain.” The Journal of Neuroscience: The Official Journal of the Society for Neuroscience 33 (50): 19434–41.

Kleiner, M., Brainard, D., & Pelli, D. 2007. “What’s new in Psychtoolbox-3?” Perception, 36, ECVP Abstract Supplement.

Kononowicz, T.W., and van Rijn, H. 2015. “Single Trial Beta Oscillations Index Time Estimation.” Neuropsychologia 75 (August): 381–89.

Kulashekhar, S., Pekkola, J., Palva, J.M., and Palva, S. 2016. “The Role of Cortical Beta Oscillations in Time Estimation.” Human Brain Mapping 37 (9): 3262–81.

Lakatos, P., Karmos, G., Mehta, A.D., Ulbert, I., and Schroeder, C.E. 2008. “Entrainment of Neuronal Oscillations as a Mechanism of Attentional Selection.” Science 320 (5872): 110–13.

Lewis, P. A., and Miall, R.C. 2003. “Brain Activation Patterns during Measurement of Sub-and Supra-Second Intervals.” Neuropsychologia 41 (12): 1583–92.

Maris, E., and Oostenveld, R. 2007. “Nonparametric Statistical Testing of EEG- and MEG-Data.” Journal of Neuroscience Methods 164 (1): 177–90.

Matell, M.S., and Meck, W.H. 2004. “Cortico-Striatal Circuits and Interval Timing: Coincidence Detection of Oscillatory Processes.” Brain Research. Cognitive Brain Research 21 (2): 139–70.

Matthews, W.J., and Meck, W.H. 2016. “Temporal Cognition: Connecting Subjective Time to Perception, Attention, and Memory.” Psychological Bulletin 142 (8): 865–907.

Oostenveld, R., Fries, P., Maris, E., and Schoffelen, J.-M. 2011. “FieldTrip: Open Source Software for Advanced Analysis of MEG, EEG, and Invasive Electrophysiological Data.” Computational Intelligence and Neuroscience 2011: 156869.

Palva, S., and Palva, J.M. 2007. “New Vistas for Alpha-Frequency Band Oscillations.”Trends in Neurosciences 30 (4): 150–58.

Pelli, D.G. 1997. “The VideoToolbox software for visual psychophysics: Transforming numbers into movies.” Spatial Vision, 10, 437–442.

Pfurtscheller, G., and Lopes da Silva, F.H. 1999. “Event-Related EEG/MEG Synchronization and Desynchronization: Basic Principles.” Clinical Neurophysiology: Official Journal of the International Federation of Clinical Neurophysiology 110 (11): 1842–57.

Rohenkohl, G., and Nobre, A.C. 2011. “Alpha Oscillations Related to Anticipatory Attention Follow Temporal Expectations.” The Journal of Neuroscience: The Official Journal of the Society for Neuroscience 31 (40). Society for Neuroscience: 14076–84.

Turek, F.W. 1985. “Circadian Neural Rhythms in Mammals.” Annual Review of Physiology 47: 49–64.

Weisz, N., Hartmann, T., Müller, N., Lorenz, I., and Obleser, J. 2011. “Alpha Rhythms in Audition: Cognitive and Clinical Perspectives.” Frontiers in Psychology 2 (April): 73.

Wilsch, A., Henry, M.J., Herrmann, B., Maess, B., and Obleser, J. 2015. “Alpha Oscillatory Dynamics Index Temporal Expectation Benefits in Working Memory.” Cerebral Cortex 25 (7): 1938–46.

Zumer, J. M., Scheeringa, R., Schoffelen, J.-M., Norris, D.G., and Jensen, O. 2014.“Occipital Alpha Activity during Stimulus Processing Gates the Information Flow to Object-Selective Cortex.” PLoS Biology 12 (10): e1001965.

